# Attempted fractionation of LB Lennox medium via reversed phase high performance liquid chromatography

**DOI:** 10.1101/2020.09.26.315150

**Authors:** Wenfa Ng

## Abstract

Mass balance analysis is a highly useful tool for chemical engineering analysis of biological processes. Specifically, composition and amounts of inputs to a cell could be correlated with measurement of metabolic byproducts and outputs for inferring metabolic fluxes flowing through specific cellular metabolic pathways. However, the composition of many common microbiological growth medium remains ill-defined with batch-to-batch variation. Thus, a need exists for developing methods for effective fractionation and separation of common growth medium. Using 5 g/L yeast extract, 10 g/L tryptone, and LB Lennox medium as model systems, this work attempted the fractionation of the three complex growth mixtures using C-18 reversed phase high performance liquid chromatography (RP-HPLC). Results revealed no effective fractionation of the three mixtures. More importantly, experiment results indicated that appropriate choice of detection wavelength for visualizing the chromatogram made a huge difference to understanding the effectiveness of fractionation achieved. Specifically, in the case of yeast extract, tryptone and LB Lennox medium, 194 nm may be a more appropriate detection wavelength compared to 280 nm. Collectively, C-18 RP-HPLC was not effective in separating 5 g/L yeast extract, 10 g/L tryptone and LB Lennox medium with hydrophilic mobile phases (ethanol/water mixture).

**Graphical abstract:** **Short description:** Visualization of the reversed phase HPLC chromatogram at 194 nm revealed broad peaks and lack of distinct peaks indicative of fractionation of LB Lennox medium by the chromatography method. More importantly, complex ensemble of different components in the growth medium present a significant separation challenge that precludes separation or fractionation by modern liquid chromatography methods. Overall, combinatorial use of both hydrophobic and hydrophilic mobile phase with a C-18 reversed phase column could reveal the presence of a couple of main fractions in a mixture otherwise unable to be separated into distinct components.

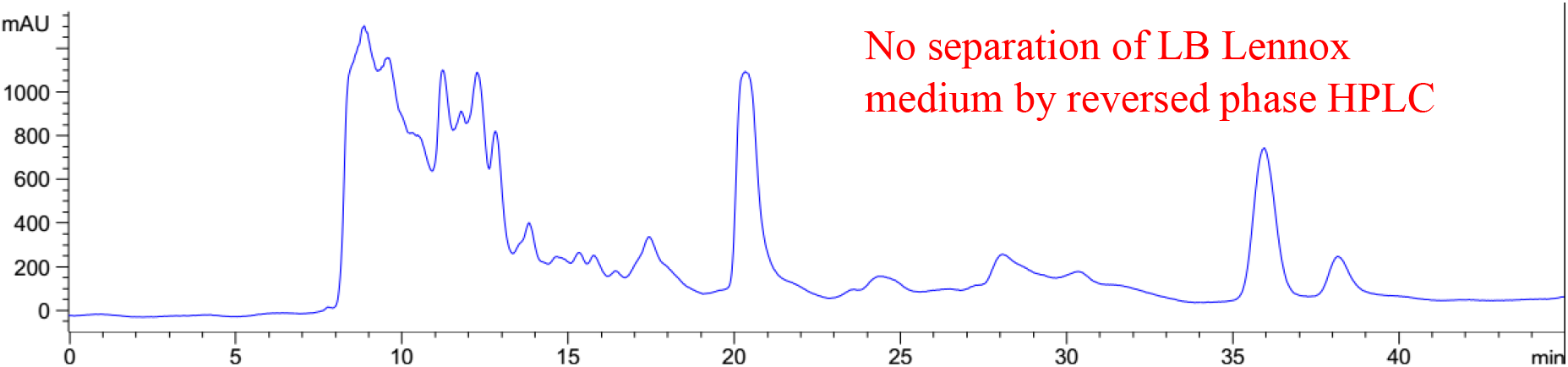

## 1. Introduction

Growth media for microbiology are complex mixture of components necessary for the cultivation of various microorganisms, especially for complex media where many of the components are not defined in substance and relative amount. Thus, use of such media for microbiology and biochemistry studies highlight one significant challenge: lack of compositional details prevents a mass balance analysis for the relative amount of each component used for growth, where such analysis could feed into whole cell metabolic models for growth.^1,2^ Additionally, with the advent of metabolomics, inability to understand the composition of growth media and that of culture broth precludes deeper insights into details concerning metabolic flux in cells along different metabolic pathways and cycles.^3–7^ Finally, batch to batch variability may exist in complex microbiological media which could impact on bacteria growth performance or product titer.^8^ But, how do we elucidate the composition of complex growth medium?

Using the common microbiological growth medium for bacteria, LB Lennox, as model system, this study attempted to separate the complex medium into different fractions through reversed phase high performance liquid chromatography (RP-HPLC). The principal objective was to understand the extent in which RP-HPLC could fractionate LB Lennox medium based on molecular weight as well as its affinity to a hydrophobic reversed phase liquid chromatography separation column. Given that the major components of LB Lennox are tryptone and yeast extract, the study first attempted the separation of the two complex undefined mixtures to help lend clarity to any problems that could be encountered in the separation of LB Lennox medium.

Experiment results revealed that the use of hydrophilic mobile phase mixture of ethanol and water did not provide C-18 RP-HPLC the ability to effectively fractionate 5 g/L yeast extract, 10 g/L tryptone and LB Lennox medium into different fractions. More importantly, the data from variable wavelength detector and diode array detector revealed that appropriate choice of detection wavelength played a critical role in visualizing the chromatogram obtained, where 194 nm provided a more informative view of the elution profile from the column in the case of 5 g/L yeast extract, 10 g/L tryptone, and LB Lennox medium. Future work could use a combination of hydrophobic and hydrophilic mobile phases in effecting a better fractionation of LB Lennox medium, 5 g/L yeast extract, and 10 g/L tryptone.

## 2. Materials and methods

### 2.1 Sample preparation

Yeast extract was purchased from Oxoid while tryptone and LB Lennox medium were purchased from Difco. They were used as is and dissolved in deionized water prior to autoclave sterilization at 121 °C for 20 minutes in glass bottles and glass shake flask. The mixtures were allowed to cool to room temperature prior to aliquot for high performance liquid chromatography analysis. 5 mL of the various mixtures were placed in 15 mL polypropylene centrifuge tubes, and the mixtures filtered through 0.22 μm nylon membrane filters prior to chromatographic analysis.

### 2.2 Reverse phase high performance liquid chromatography (RP-HPLC) analysis

10 μL of filtered mixture was injected into the column of Agilent 1200 series high performance liquid chromatography (HPLC) instrument, with Poroshell 120 SB-C18 as column. Sufficient time was allowed for the column to equilibrate to temperature and flow rate settings prior to commencement of analysis. Mobile phase chosen was a mixture of ethanol and water in isocratic elution profile. Both variable wavelength detector and diode array detector were used for visualizing the chromatogram, where wavelength used for variable wavelength detector was 280 nm, and the wavelength set for diode array detector was 280, 264, 254, 194 and 380 nm with 360 nm as reference.

## 3. Results and Discussion

Reversed phase high performance liquid chromatography (RP-HPLC) was first used in separating one of the major components of LB Lennox medium, 5 g/L yeast extract. With an injection volume of 10 μL, several fractions were observed eluting out of the HPLC column, each with distinct peaks on the chromatogram, but separation of the mixture into various components such as amino acids, proteins, lipids, was not achieved (Figure 1).

**Figure 1:**
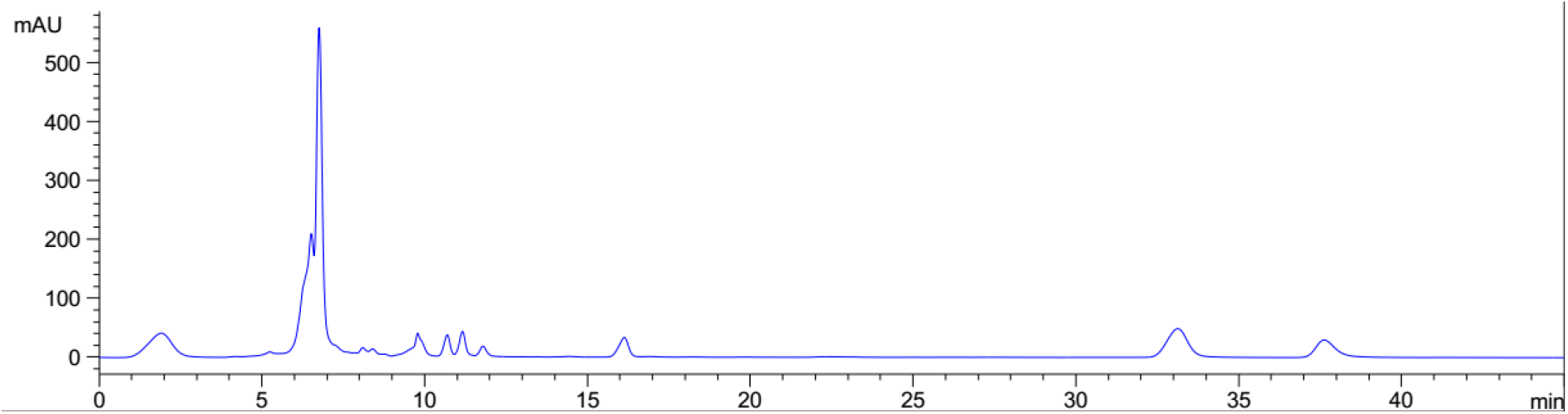
Fractionation of diluted 5 g/L yeast extract with reversed phase high performance liquid chromatography (RP-HPLC) into several fractions. Detection by variable wavelength detector at 280 nm, Column: Agilent Technologies Poroshell 120 SB-C18, Flow rate = 0.2 ml/min, 5/95 Ethanol/water (%/%) mobile phase for isocratic elution, 25 °C, 81 bar pressure, Injection volume = 10 μL

As yeast extract is a mixture of hydrophobic and hydrophilic components, reduction in flow rate from 0.6 ml/min to 0.3 ml/min increased the residence time of the compounds in yeast extract in the column (Figure S1 and S2). Changes of isocratic eluent from 50/50 (%/%) ethanol/water to 10/90 (%/%) ethanol/water increases the polarity of the mobile phase, which favours the desorption of yeast extract components from the C-18 reversed phase column. This manifested as broader peaks at the same retention time in the chromatogram (Figure S3). On the other hand, use of an 80/20 (%/%) ethanol/water eluent compared to 50/50 (%/%) ethanol/water eluent increased the hydrophobicity of the mobile phase, which favours adsorption of components to the reversed phase column. (Figure S4). Thus, given the presence of both hydrophobic and hydrophilic components in yeast extract, a hydrophilic mobile phase should be used to help desorb hydrophilic components of yeast extract into different fractions. To this end, a 5/95 (%/%) ethanol/water mixture was used as mobile phase, which helps to fractionate 5 g/L yeast extract into more fractions (Figure S5). With the goal of reducing flow rate for increasing contact time between mobile phase and column, thereby, aiding in gaining a more granular separation of the complex mixture, a slower flow rate of 0.1 ml/min was used (Figure S6). However, a slower flow rate did not enable the fractionation of more components; thus, highlighting that a different strategy is needed to elute the more hydrophobic components which remain adsorbed on the C-18 reversed phase column.

Attempts at fractionating 10 g/L tryptone, a component of LB Lennox growth medium also did not succeed. Specifically, a single peak eluted out of RP-HPLC under isocratic elution conditions (Figure 2). A second run of the analysis reveals the same phenomenon of a distinct peak at 5 minutes retention time (Figure S7).

**Figure 2:**
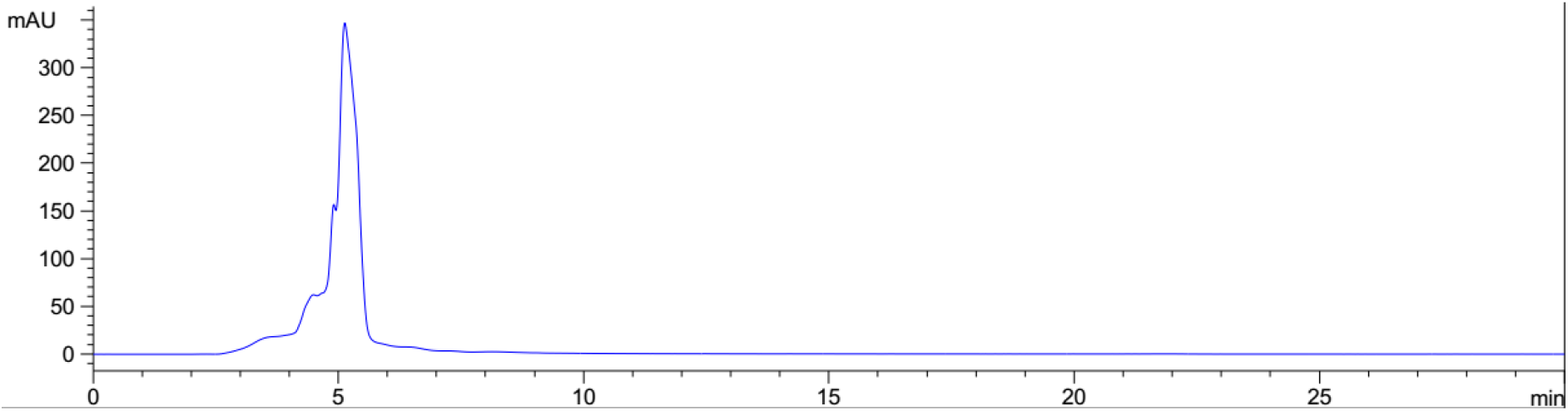
Inability to fractionate 10 g/L Tryptone into different fractions using reversed phase high performance liquid chromatography. Detection by variable wavelength detector at 280 nm, Column: Agilent Technologies Poroshell 120 SB-C18, Flow rate = 0.3 ml/min, 80/20 Ethanol/water (%/%) mobile phase for isocratic elution, 35 °C, 194 bar pressure, Injection volume = 10 μL

Using the set of detection wavelengths available to a diode array detector, attempts were made to visualize the different fractions of 5 g/L yeast extract separated by RP-HPLC under characteristic detection wavelength of specific functional groups in organic compounds. It could be seen in Figure 3 that similar chromatograms were obtained for detection wavelength of 280, 264 and 254 nm. However, the peaks of the same chromatogram appeared to be broader under a detection wavelength of 194 nm. Finally, 380 nm detection wavelength did not detect peaks of appreciable intensity.

**Figure 3:**
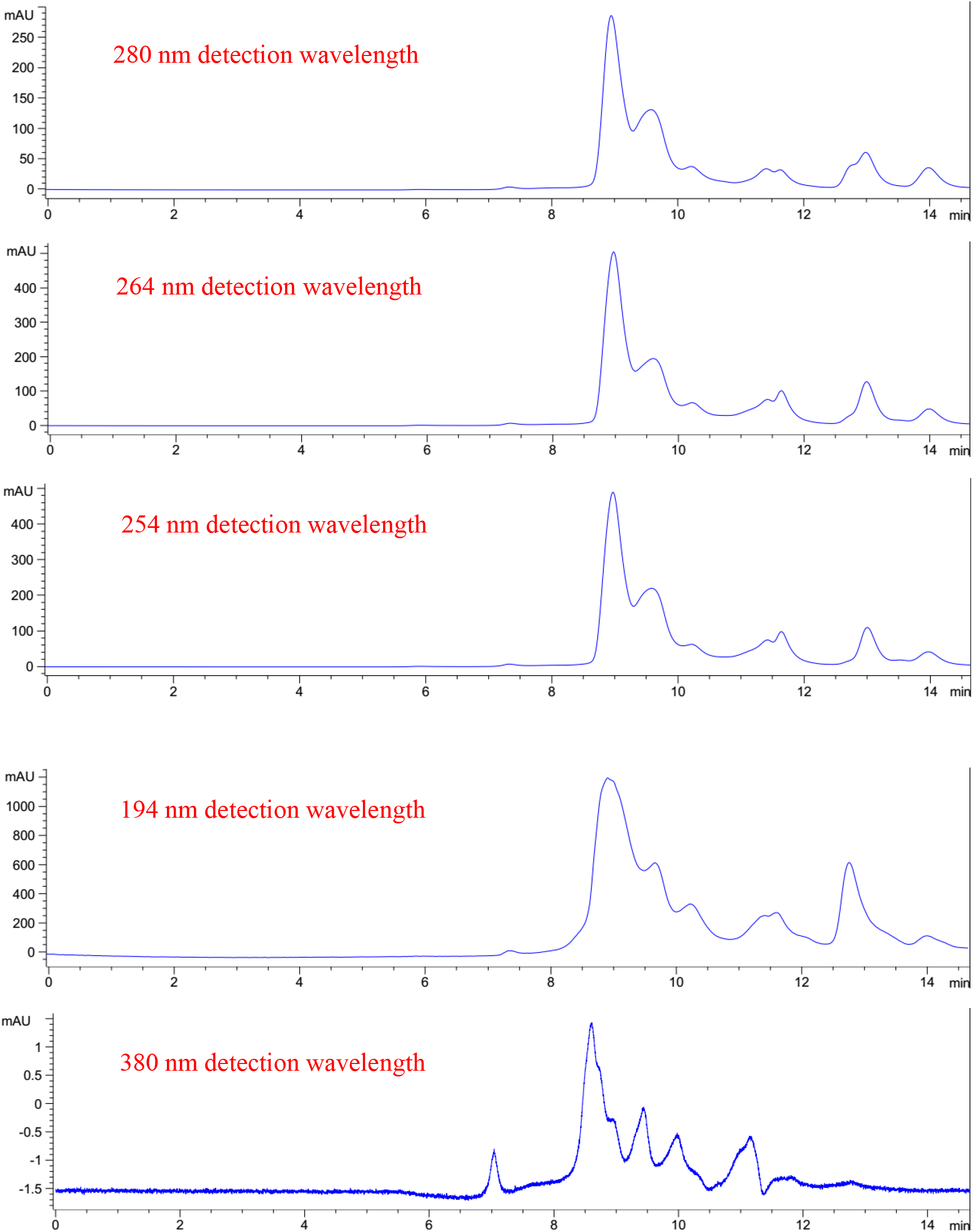
Attempted fractionation of 5 g/L yeast extract. Column: Agilent Technologies Poroshell 120 SB-C18, Flow rate = 0.2 ml/min, 5/95 Ethanol/water (%/%) mobile phase for isocratic elution, 25 °C, 70 bar pressure, Injection volume = 10 μL

Similarly, attempts were also made to visualize the different fractions that elute out of the C-18 reversed phase column during the fractionation of 10 g/L tryptone (Figure 4). Readout at different wavelengths of a diode array detector revealed that similar elution profile was observed in the chromatogram for 280, 264, and 254 nm. Specifically, broader peaks were observed for chromatogram visualized at 254 and 264 nm compared to 280 nm in the retention time region between 5 and 25 minutes, while the sharp peak at retention time of 36 mins was consistent for all three visualization wavelengths of 254, 264 and 280 nm. This indicated that many components of tryptone were hydrophilic and eluted early during chromatography. A similar chromatogram was also visualized at 194 nm, but with much broader and intense peaks compared to those visualized at 254 and 264 nm. Similar to the case for 5 g/L yeast extract, 380 nm detection wavelength did not detect peaks of appreciable intensity.

**Figure 4:**
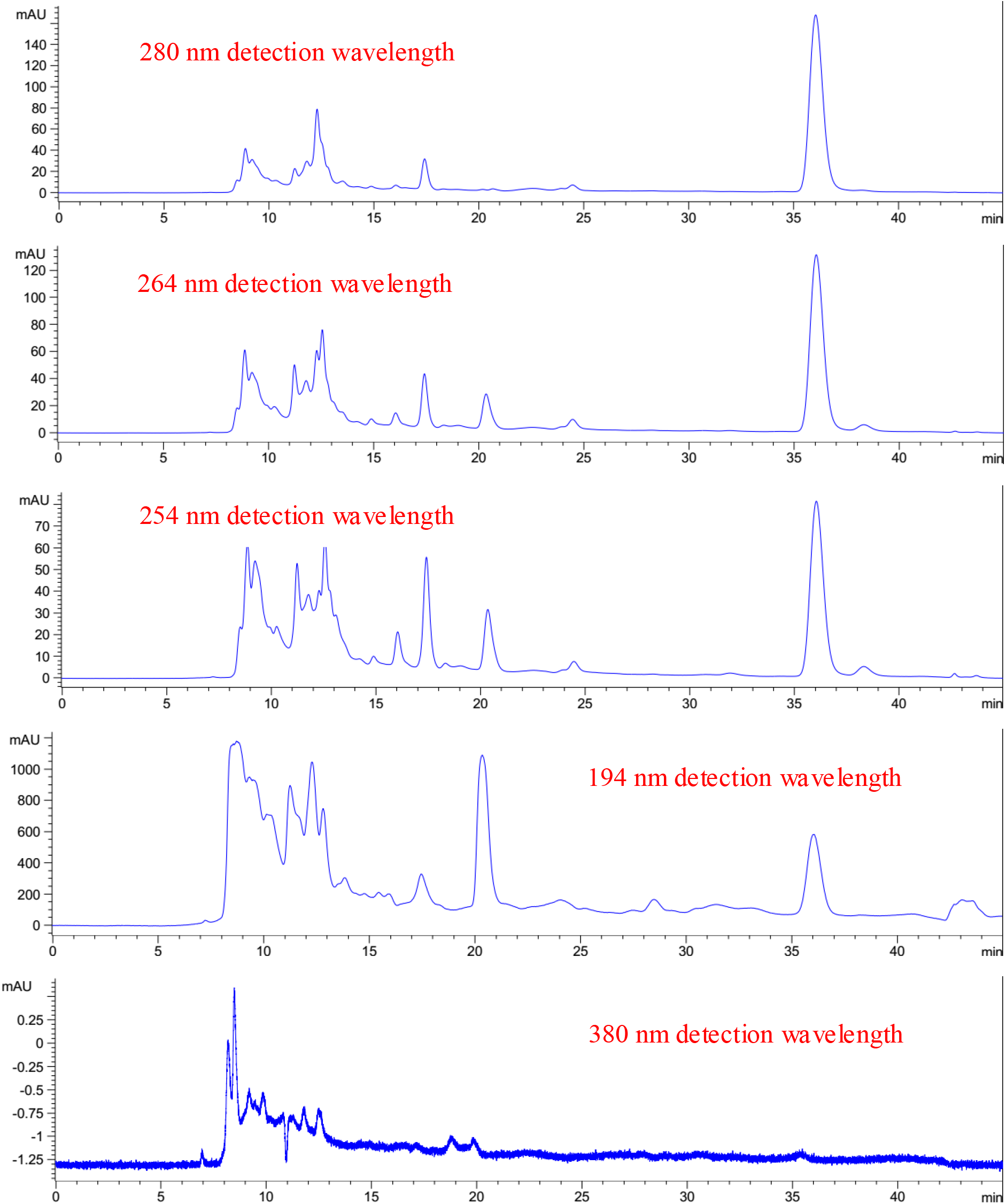
Attempted fractionation of 10 g/L Tryptone. Column: Agilent Technologies Poroshell 120 SB-C18, Flow rate = 0.2 ml/min, 5/95 Ethanol/water (%/%) mobile phase for isocratic elution, 25 °C, 71 bar pressure, Injection volume = 10 μL

LB Lennox medium was subjected to fractionation by reversed phase high performance liquid chromatography (RP-HPLC), but the complexity of the mixture does not allow adequate fractionation. Figure 5 reveals commingled peaks eluting from a C-18 reversed phase column loaded with undiluted LB Lennox medium. The chromatogram was visualized at different wavelengths. Due to the small difference in visualization wavelengths, similar chromatograms were obtained at 280, 264 and 254 nm, with those visualized at 264 and 254 nm showing a higher degree of similarity than those at 280 nm. However, broader and more intense peaks were visualized at 194 nm. Finally, no appreciable peaks were visualized at 380 nm detection wavelength.

**Figure 5:**
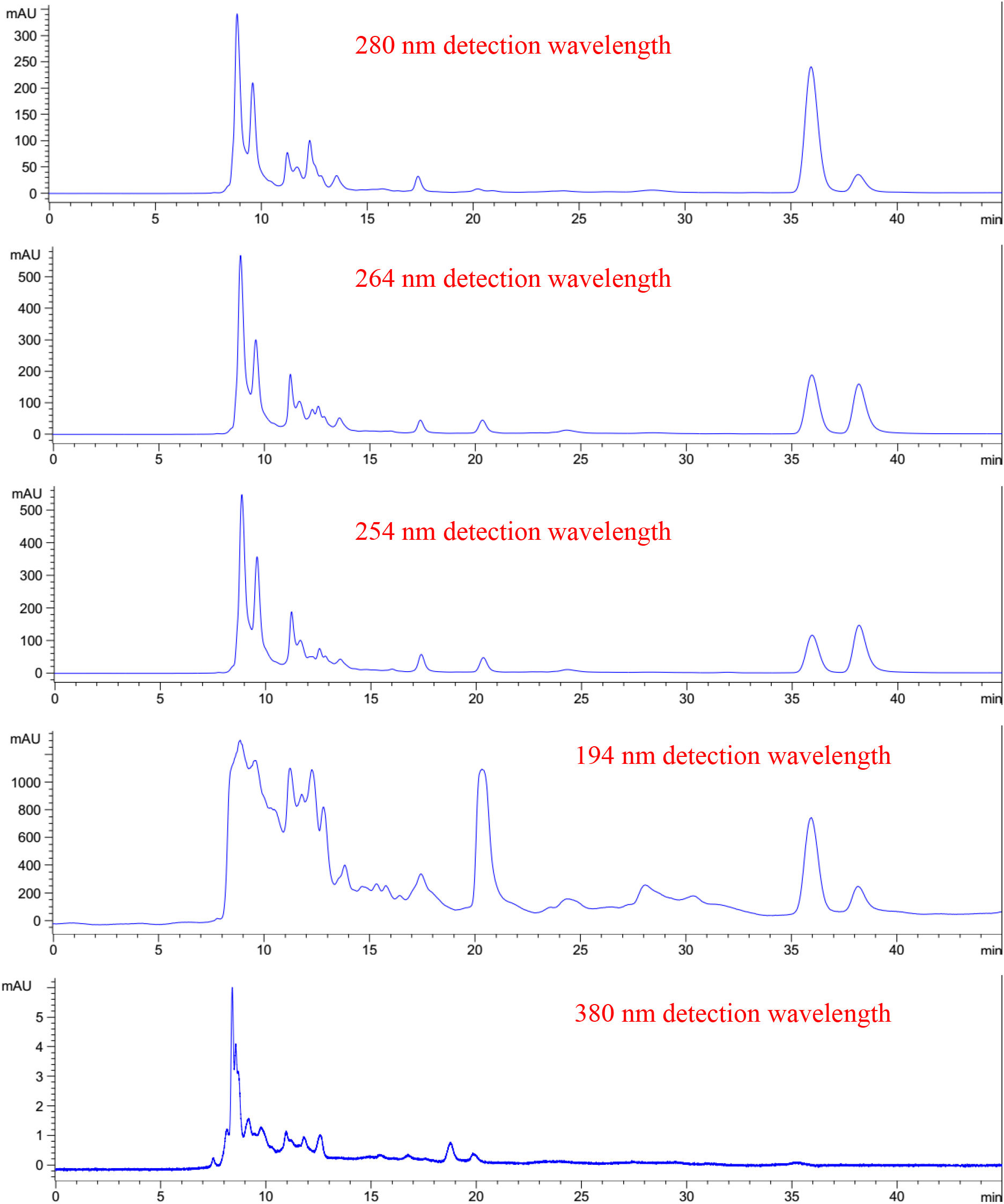
Attempted fractionation of LB Lennox medium. Column: Agilent Technologies Poroshell 120 SB-C18, Flow rate = 0.2 ml/min, 5/95 Ethanol/water (%/%) mobile phase for isocratic elution, 25 °C, 71 bar pressure, Injection volume = 10 μL

Data from diode array detector analysis of elution profile at different wavelengths revealed that there was no effective fractionation of 5 g/L yeast extract, 10 g/L tryptone and LB Lennox medium by reversed phase high performance liquid chromatography. More importantly, the data highlighted the importance of using the appropriate detection wavelength for understanding the elution properties of the adsorbate from the reversed phase column. For example, while 280 nm detection wavelength revealed poor separation of different components in the mixture, detection at 194 nm revealed even broader peaks. However, for the case of understanding the fractionation of components of LB Lennox medium, such as yeast extract and tryptone, 194 nm might be the more appropriate wavelength for understanding how different elution conditions affect the separation of different components in complex mixture such as yeast extract, tryptone and LB Lennox medium.

## 4. Conclusions

Compositional analysis of growth medium provides important information on the relative abundance of different nutrients, which can be fed into a mass balance analysis of cellular processes common in simulation models of cell metabolism. However, given the widespread use of complex medium in microbiology and biotechnology, and the tremendous difficulty of gaining an understanding of the different constituents of such growth media, significant differences exist between different models offered for understanding cellular metabolism. Specifically, while metabolomics studies in combination with metabolic engineering simulation could provide some estimates of the relative flux between different metabolic pathways, significant variation in estimates exist between different models for the same species under identical conditions. Motivated by the curiosity to understand the composition of LB Lennox, yeast extract and tryptone, at least at the level of how many fractions comprise the growth medium, reversed phase high performance liquid chromatography was used in separating the three complex growth mixtures. Exploration of the parameter space for separating 5 g/L yeast extract revealed that a combination of hydrophobic and hydrophilic mobile phases was needed for more effective fractionation of the complex mixture. On the other hand, data from diode array detector monitoring of separation performance at different wavelengths revealed that appropriate choice of detector wavelength for visualizing the different fractions eluting from the column was critical for accurate understanding of the fractionation of the mixture. Overall, 5 g/L yeast extract, 10 g/L tryptone and LB Lennox medium were not effectively fractionated by C-18 reversed phase high performance liquid chromatography with hydrophilic (ethanol/water) mobile phase, but combined use of both hydrophilic and hydrophobic mobile phases might effect better fractionation of the complex growth media in future studies.

## Conflicts of interest

The author declares no conflicts of interest.

## Funding

The author thank the National University of Singapore for financial support.

## Notes

### Competing Interest Statement

The authors have declared no competing interest.

