## Supplementary materials for "Attempted fractionation of LB Lennox medium via reversed phase high performance liquid chromatography"

Wenfa Ng

Department of Chemical and Biomolecular Engineering, National University of Singapore

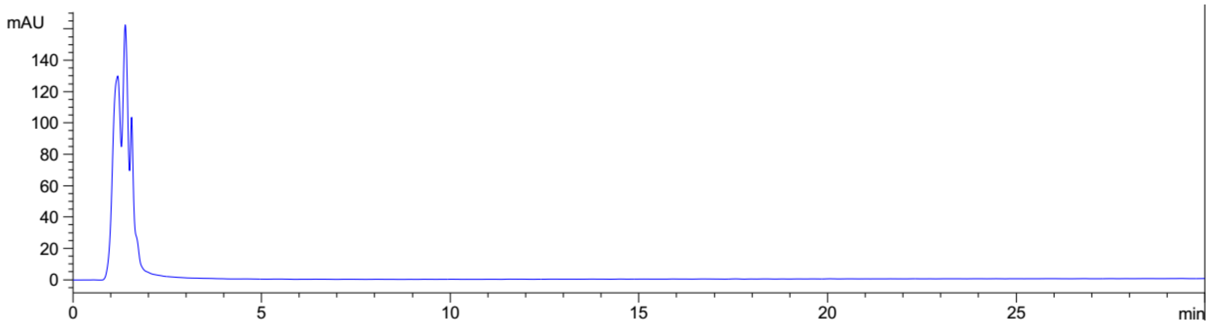


**Figure S1:** Attempted fractionation of 5 g/L yeast extract with reverse phase high performance liquid chromatography (RP-HPLC). Detection by variable wavelength detector at 280 nm, Column: Agilent Technologies Poroshell 120 SB-C18, Flow rate = 0.6 ml/min, 50/50 Ethanol/water (%/%) mobile phase for isocratic elution, 35 $^{o}$C, 240 bar pressure, Injection volume = 10 µL

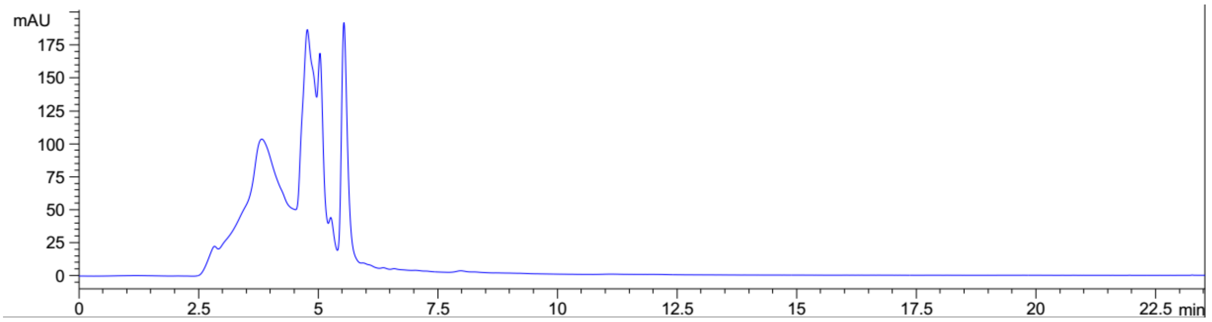


**Figure S2:** Attempted fractionation of 5 g/L yeast extract with RP-HPLC. Detection by variable wavelength detector at 280 nm, Column: Agilent Technologies Poroshell 120 SB-C18, Flow rate = 0.3 ml/min, 50/50 Ethanol/water (%/%) mobile phase for isocratic elution, 35 $^{o}$C, 240 bar pressure, Injection volume = 10 µL

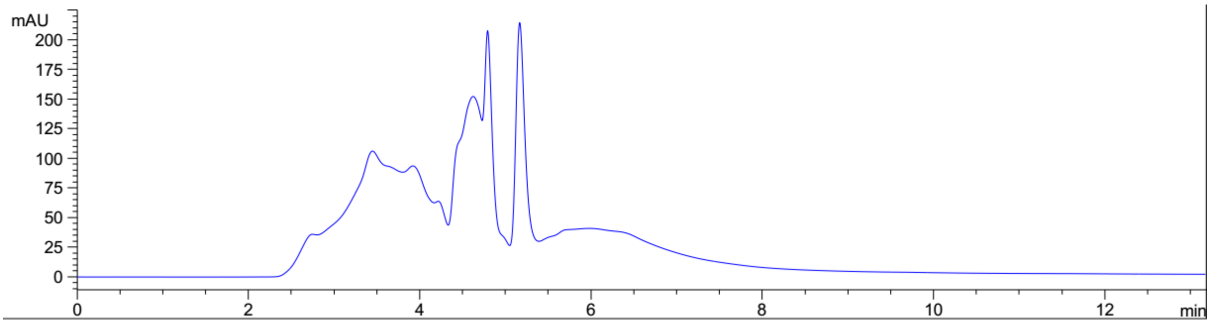


**Figure S3:** Attempted fractionation of 5 g/L yeast extract with RP-HPLC. Detection by variable wavelength detector at 280 nm, Column: Agilent Technologies Poroshell 120 SB-C18, Flow rate = 0.3 ml/min, 10/90 Ethanol/water (%/%) mobile phase for isocratic elution, 35 $^{o}$C, 126 bar pressure, Injection volume = 10 µL

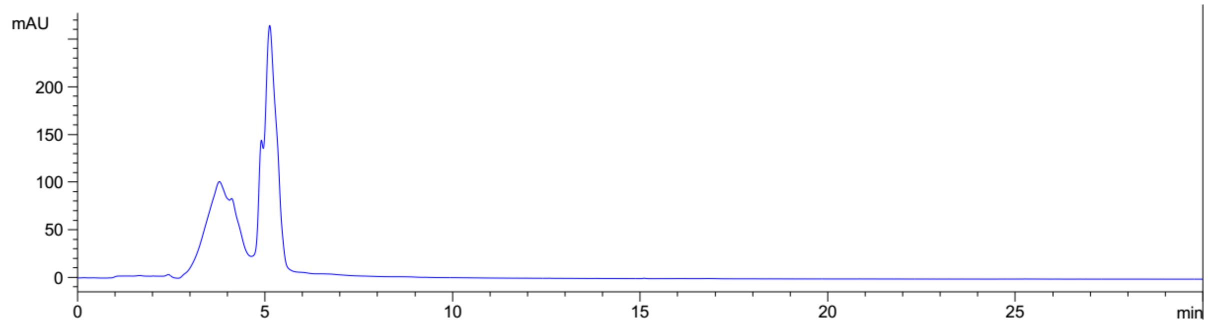


**Figure S4:** Attempted fractionation of 5 g/L yeast extract with RP-HPLC. Detection by variable wavelength detector at 280 nm, Column: Agilent Technologies Poroshell 120 SB-C18, Flow rate = 0.3 ml/min, 80/20 Ethanol/water (%/%) mobile phase for isocratic elution, 35 $^{o}$C, 196 bar pressure, Injection volume = 10 µL

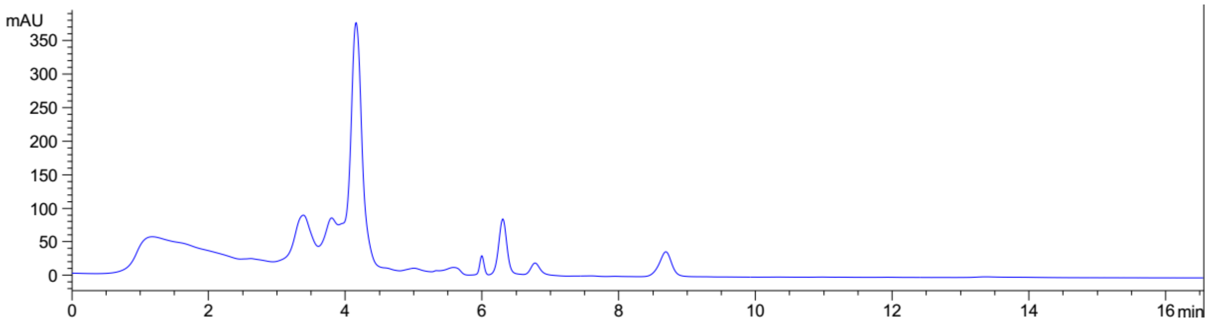


**Figure S5:** Attempted fractionation of 5 g/L yeast extract with RP-HPLC. Detection by variable wavelength detector at 280 nm, Column: Agilent Technologies Poroshell 120 SB-C18, Flow rate = 0.3 ml/min, 5/95 Ethanol/water (%/%) mobile phase for isocratic elution, 25 $^{o}$C, 121 bar pressure, Injection volume = 10 µL

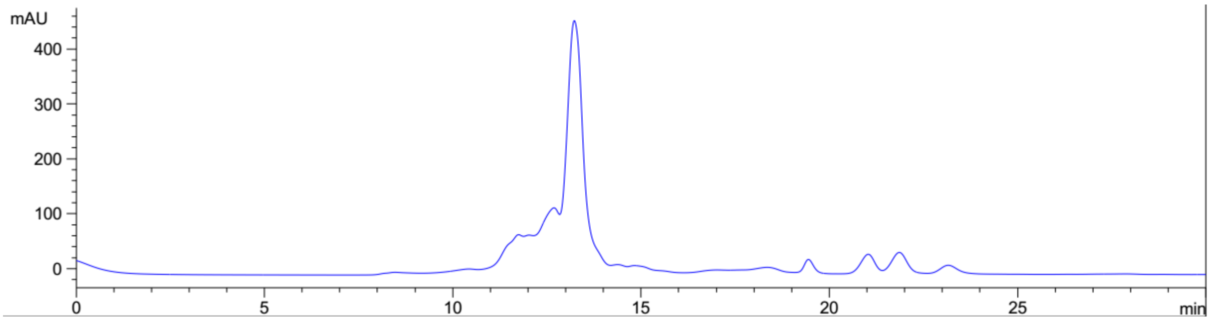


**Figure S6:** Attempted fractionation of 5 g/L yeast extract with RP-HPLC. Detection by variable wavelength detector at 280 nm, Column: Agilent Technologies Poroshell 120 SB-C18, Flow rate = 0.1 ml/min, 5/95 Ethanol/water (%/%) mobile phase for isocratic elution, 25 $^{o}$C, 41 bar pressure, Injection volume = 10 µL

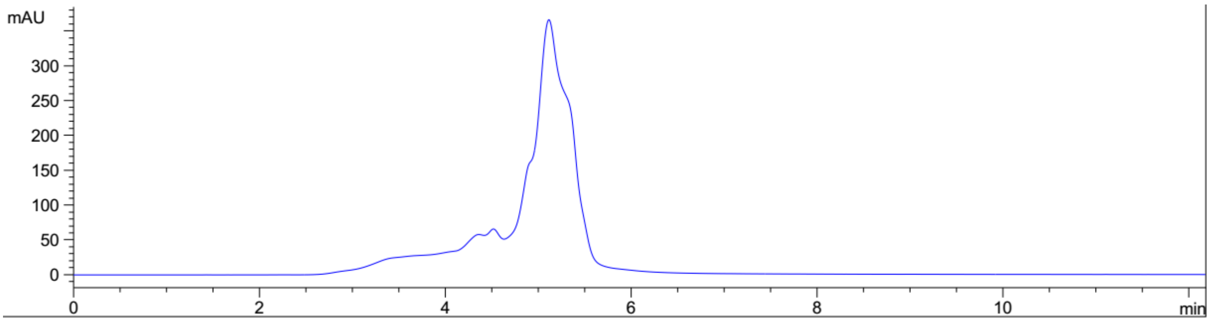


**Figure S7:** Inability to fractionate 10 g/L Tryptone into different fractions using reversed phase high performance liquid chromatography (RP-HPLC). Detection by variable wavelength detector at 280 nm, Column: Agilent Technologies Poroshell 120 SB-C18, Flow rate = 0.3 ml/min, 80/20 Ethanol/water (%/%) mobile phase for isocratic elution, 35 $^{o}$C, 194 bar pressure, Injection volume = 10 µL

### Conflicts of interest

The author declares no conflicts of interest.

### Funding

The author thank the National University of Singapore for financial support.
